# Engineering a pH-sensitive humanized infliximab with improved potency and developability using STEM™

**DOI:** 10.64898/2026.08.18.745573

**Authors:** Paul D Entzminger, Kevin C Entzminger, Jonathan K Fleming, Alex Samadi, Lisa Yuko Espinosa, Yuko Hiramoto, Shigeru CJ Okumura, Toshiaki Maruyama

## Abstract

**Background:** Tumor necrosis factor-α inhibitors such as infliximab and adalimumab have transformed autoimmune disease treatment; however, infliximab is a mouse-human chimeric antibody that remains immunogenic, is associated with self-association/aggregation liability, and requires prolonged intravenous administration. We humanized infliximab and engineered infliximab-derived candidates with improved potency and developability.

**Methods:** Infliximab complementarity-determining regions were grafted onto human germline frameworks to generate humanized infliximab. STage-Enhanced Maturation (STEM^TM^) technology produced an affinity-matured clone (hInBG4), followed by targeted amino-acid substitutions in the complementarity-determining regions to generate LW2Y, LW2YR2S, and LW2YHR1K. Variants were evaluated by a cell-based tumor necrosis factor alpha neutralization assay, affinity-capture self-interaction nanoparticle spectroscopy, a baculovirus particle enzyme-linked immunosorbent assay, size-exclusion high-performance liquid chromatography, transient expression in human embryonic kidney 293 cells, and tumor necrosis factor alpha binding kinetics by biolayer interferometry, including dissociation at pH 7.4 and 5.8.

**Results:** All three variants showed two-to three-fold higher neutralization potency than chimeric infliximab and outperformed adalimumab. Affinity-capture self-interaction nanoparticle spectroscopy shifts decreased from double-digit parental values to low single digits, while baculovirus particle binding ratios remained acceptable. Size-exclusion chromatography showed cleaner monomer peaks with reduced tailing, and expression increased relative to humanized infliximab. LW2Y combined very high affinity at pH 7.4 with markedly faster dissociation at pH 5.8, consistent with pH-dependent antigen release.

**Conclusions:** Humanization, affinity maturation, and targeted complementarity-determining region re-engineering generated infliximab-derived candidates with improved potency and developability and identified LW2Y as a lead for further preclinical evaluation.

**Statement of Significance:** STEM™ engineering produced a pH-sensitive, humanized infliximab lead with improved potency and developability. LW2Y shows two-to three-fold stronger TNF-α neutralization, low self-association, improved expression, and accelerated antigen dissociation at pH 5.8, supporting advancement into preclinical pharmacokinetic and efficacy studies.

## INTRODUCTION

Monoclonal antibodies that neutralize tumor necrosis factor-α (TNF-α) have become a mainstay in the treatment of rheumatoid arthritis, inflammatory bowel disease, and other immune-mediated disorders. Several anti-TNF-α agents are approved, including the chimeric mouse–human antibody infliximab and the fully human antibody adalimumab [1]. While these drugs provide substantial clinical benefit, their long-term use can be limited by immunogenicity and secondary loss of response due to anti-drug antibodies [1, 2]. Infliximab, in particular, is also associated with aggregation and self-association liabilities and the need for prolonged intravenous infusions, which complicate administration, formulation, and manufacturing. Consistent with these observations, structural and solution studies have shown that infliximab can undergo reversible self-association mediated by Fab-Fab interactions, providing a mechanistic rationale for developability-focused optimization of this clinically validated scaffold [3].

Infliximab is composed of murine variable regions fused to human immunoglobulin G1 (IgG1) constant domains and therefore contains a substantial proportion of murine sequence [1]. The murine variable domain is a major driver of human anti-chimeric antibody responses, which can accelerate clearance, reduce efficacy, and increase the risk of infusion reactions [1, 2]. Concomitant administration of immunomodulators such as methotrexate mitigates but does not abolish infliximab immunogenicity in patients with Crohn’s disease and related indications [2]. In contrast, humanized and fully human anti-TNF-α antibodies generally exhibit lower anti-drug antibody rates, although they can still be immunogenic [1]. These considerations motivate the development of “biobetter” anti-TNF-α antibodies that combine reduced immunogenicity risk with improved biophysical properties and patient convenience.

Beyond immunogenicity, the developability of a therapeutic antibody has emerged as a critical determinant of clinical success [4–6]. High-throughput assays such as affinity-capture self-interaction nanoparticle spectroscopy (AC-SINS) and baculovirus particle enzyme-linked immunosorbent assay (BVP ELISA) are widely used to detect self-association and polyspecificity early in discovery. AC-SINS detects weak antibody self-interaction by measuring plasmon peak shifts of antibody-decorated gold nanoparticles, and increased AC-SINS wavelength shift Δλmax (nm) correlates with aggregation liability, high viscosity, and poor colloidal stability [7–10]. BVP ELISA uses baculovirus particles as a multivalent, heterogeneous surface to quantify non-specific binding; high BVP ratios have been associated with faster in vivo clearance and with antibodies that exhibit increased polyspecificity in other assays [11–14]. Together with size-exclusion chromatography (SE-HPLC), expression titer, and sequence liability assessment, these assays help triage candidates with unfavorable developability profiles and enrich for molecules with “drug-like” specificity [4–6, 10, 13, 15].

In addition to humanization and developability optimization, a complementary strategy to improve potency is to engineer pH-dependent antigen binding. In this paradigm, the antibody binds antigen tightly at neutral pH but releases it in the mildly acidic endosome, enabling FcRn-mediated antibody salvage while promoting antigen degradation and thus increasing potency [16]. This strategy has been used for TNF-α-neutralizing mAbs, including one monovalent, pH-sensitive anti-TNF-α antibody designed to reduce immunogenicity in vivo and another engineered using a histidine-scanning approach that introduced pH dependence but in some cases reduced neutral-pH binding strength [17, 18]. Collectively, these studies support the feasibility of engineering pH-dependent anti-TNF-α antibodies while underscoring the value of approaches that preserve (or improve) neutral-pH potency and concurrently address developability.

We previously developed the STage-Enhanced Maturation (STEM™) technology, which combines rational complementarity-determining region (CDR) design with iterative selection to improve both affinity and developability of antibody leads [19, 20]. In the present study, we applied STEM™ to the clinically validated but immunogenic anti-TNF-α antibody infliximab to generate a humanized scaffold and a panel of engineered variants. We show that these infliximab-derived antibodies exhibit enhanced TNF-α neutralization potency, improved self-association and non-specific binding profiles, and, most notably in the case of LW2Y, pronounced pH-dependent antigen dissociation, identifying LW2Y as a promising biobetter lead for further preclinical evaluation.

## MATERIALS AND METHODS

### Antibody design, humanization and cloning

The amino acid sequences of the chimeric anti-TNF-α antibody infliximab (Remicade®) and the fully human antibody adalimumab (Humira®) were obtained from public patent and sequence databases. The variable heavy (VH) and light (VL) region sequences of infliximab were aligned to human germline genes using IgBLAST (NCBI), and IGKV6-21 and IGHV3-72 were selected as human framework acceptor sequences based on highest sequence identity and preservation of canonical CDR loop structures. Mouse CDRs were grafted onto the selected human germline frameworks. Humanized infliximab (hInfliximab) variable-and constant-region genes were synthesized (Twist Bioscience). For CDR diversification using the STage-Enhanced Maturation (STEM™) platform, focused libraries were designed at preselected CDR positions based on amino acid usage and positional frequencies observed in human antibodies to further bias the sequences toward human-like profiles. Degenerate codons (e.g. NNK or tailored wobbled base mixes) were used to maximize amino acid diversity while keeping total library complexity within the transformation capacity of electroporated E. coli ∼1 × 10¹⁰ colony-forming units (cfu). VH-and VL-genes of selected clones and those of infliximab, hInfliximab, and adalimumab were cloned into human IgG1 κ expression vectors using standard restriction enzyme digestion and ligation cloning. Heavy-and light-chain expression cassettes were placed under separate cytomegalovirus promoters, and the constructs included a human IgG1 constant region with an unmodified Fc. All plasmids were confirmed by Sanger sequencing prior to expression.

### Phage panning

Phage panning of STEM™ libraries was performed as described previously [19]. In Stage 1, six single CDR mutant libraries were panned on 1 µg/mL of recombinant TNF-α to remove non-productive clones and to enrich all possible CDR mutants that retain binding to TNF-α (1 hour incubation time and Dulbecco’s phosphate-buffered saline (PBS)-washes 3 times for round 1, 5 washes for round 2-3, and 10 washes for round 4 with 5 min incubation time). Library DNAs were transformed into XL1-Blue cells (Monserate Biotechnology Group) for overnight growth and phage production following addition of M13KO7 Helper Phage (New England Biolabs). Phage were precipitated the following day from culture supernatant by addition of 4% PEG-8000/3% NaCl on ice, followed by pelleting and resuspension in 0.1% BSA/PBS (blocking buffer). The library phage was first transiently heated (68°C for stage 1, 71°C for Stage 2, and 72°C for 10 min for Stage 3), then cooled to improve thermoresistance as described previously [21], followed by subtraction of the library on polyspecificity reagents (PSRs) - BVP, dsDNA, ovalbumin, HSP90 fused to human IgG Fc (HSP90-hFc), and KLH-coated microtiter wells to remove polyreactive clones before selection on TNF-α. High-bind microtiter wells (Immulon 4HBX) were coated with recombinant TNF-α overnight, washed with PBS, and blocked with blocking buffer for 1 hour at 37°C. Phage were added for incubation at 37°C for 1 hour, followed by washing with PBS, elution with 0.1 M HCl, and neutralization with 2 M Tris. Neutralized phage were used to infect ER2738 cells (New England Biolabs) for overnight propagation and precipitation as described above. Panning was performed for four rounds, with increasing washing stringency in later rounds to enrich specific binders. In Stage 2, round four output phage from Stage 1 were used to amplify targeted CDR regions and combined by overlap PCR into separate light chain (Lx; paired with the wild type heavy chain) or heavy chain (Hx; paired with the wild type light chain) libraries. The Lx library and Hx libraries were panned on recombinant TNF-α (0.2 µg/mL for round 1-2, 0.1 µg/mL for round 3, and 0.05 µg/mL for round 4 and PBS-washes were 5 times for round 1-2 and 10 times for round 3 and 4 with 5 min incubation time). In Stage 3, round four output phage from Stage 2 were used to amplify light and heavy chain libraries for sequential cloning and pairing to create a single combined library. This library was panned stringently (incubation time was 1 min and PBS-washes were 5 times for round 1-3 and 10 times for round 4 with 5 min incubation time) on 100 ng/mL (round 1 and 2) and 50 ng/mL (round 3 and 4) of recombinant TNF-α for four rounds.

### Fab blocking assay

Following the final round of phage panning, single colonies were prepared from the eluted phage. Colonies were picked to 96-well culture plates with 1 mL of Super Broth medium and 50 μg/mL of carbenicillin. The culture plates were incubated at 37°C and 1 mM isopropyl ß-D-1- thiogalactopyranoside was added to induce overnight soluble Fab production at 30°C. Microtiter wells were coated with 100 μL of recombinant TNF-α receptor fused to human IgG Fc (TNFR-Fc) at 2 μg/mL in PBS. The culture supernatants containing soluble Fab prepared as above for Fab binding ELISA were mixed 1:1 with a recombinant TNF-α with 6xHis tag (0.04 µg/mL) at room temperature for 1 hour. Following washing and blocking, culture supernatants and TNF-α mixture were added to the wells for incubation at room temperature for 1 hour. After washing, bound TNF-α was detected with peroxidase conjugated rabbit anti-His antibody (Jackson ImmunoResearch) at room temperature for 1 hour. Blank well was used as a positive control and the signals were read at OD450 nm. The % reduction of signals was calculated using blank control well as 100% signal. The clones that showed blocking of TNF-α binding to TNFR-Fc were Polymerase Chain Reaction (PCR) amplified and sequenced by Sanger sequencing (Eton Biosciences).

### Crude Fab polyspecificity evaluation by ELISA

For crude Fab polyspecificity evaluation by ELISA, we routinely use HSP90-hFc and double-stranded deoxynucleic acid (dsDNA), because HSP90-hFc can be produced in large quantities at lower cost and in our experience, BVP ELISA ratios had good correlation with HSP90-hFc ELISA ratios. Microtiter wells were coated with HSP90-hFc (10 µg/mL) or dsDNA (2 µg/mL) in PBS at 4°C overnight. Culture supernatants containing Fab were mixed with peroxidase conjugated goat anti-human IgG F(ab’)_2_ specific antibody (Jackson ImmunoResearch) for 30 minutes at room temperature to form a bivalent (pseudo-IgG) complex. Following washing and blocking, culture supernatant mixtures were added to the wells for incubation at 4°C overnight. After washing, bound human Fab was detected after development with 3,3’, 5,5’-tetramethylbenzidine substrate (TMB). The reaction was stopped with 0.6 N H₂SO₄, and absorbance was read at 450 nm. The ratio was calculated by dividing the signal of each clone by the signal of blank control (0.1% BSA/PBS). Crude Fab preparations of hInfliximab and adalimumab were also included as controls in this study.

### Crude Fab polyspecificity evaluation by flow cytometry

For crude Fab polyspecificity evaluation by flow cytometry, streptavidin beads coated with biotinylated HSP90-hFc were incubated with culture supernatants containing soluble Fabs. Beads were washed with Flow Cytometry Staining Buffer (FACS) and bound Fab was detected with AlexaFluor 647-conjugated goat anti-human IgG F(ab’)_2_ specific antibody (Jackson ImmunoResearch). The ratio was calculated by dividing the signal of each clone by the signal of blank control (FACS only). Crude Fab preparations of hInfliximab and adalimumab were also included as controls in this study.

### Sequence-guided liability assessment and targeted amino-acid substitutions

The VH and VL amino-acid sequences of hInBG4 were evaluated for CDR-localized physicochemical features associated with unfavorable antibody developability. Candidate sites included aromatic or hydrophobic residues, particularly Trp and Phe, that could contribute to local hydrophobic patches and antibody self-association, as well as Arg residues that could contribute to positively charged patches and non-specific or polyreactive binding. Because the effects of individual residues depend on their sequence and structural context, these features were treated as potential risk indicators rather than definitive liabilities [22–24].

Selected hInBG4 residues were either reverted to the corresponding residue in parental infliximab, where applicable, or replaced with a conservative alternative intended to reduce local hydrophobicity or charge-related interactions while minimizing disruption of antigen binding. Substitutions were introduced individually or in combination to generate the following variants: LW2Y (VL W50Y); LW2YR2S (VL W50Y/R54S); HR1K (VH R31K); HF1Y (VH F33Y); HF3T (VH F100T); LW2YHR1K (VL W50Y and VH R31K); LW2YHF1Y (VL W50Y and VH F33Y); LW2YHF3T (VL W50Y and VH F100T); LW2YR2SHR1K (VL W50Y/R54S and VH R31K); LW2YR2SHF1Y (VL W50Y/R54S and VH F33Y); and LW2YR2SHF3T (VL W50Y/R54S and VH F100T). Residue positions are reported according to the Kabat numbering scheme [25].

### Antibody expression and purification

Antibodies were expressed by transient transfection of human embryonic kidney 293 (HEK293) cells. Briefly, suspension-adapted HEK293 cells were maintained in chemically defined serum-free medium (FreeStyle™ F17 Expression Medium supplemented with 4 mM L-glutamine and 0.1% Kolliphor P-188) at 37°C, 5-8% CO₂, with orbital shaking at 150 rpm. Cells were seeded at least one day before transfection with transfection day target of ∼2.5 × 10⁶ cells/mL. A heavy-and light-chain bicistronic plasmid was transfected using a polyethylenimine-based reagent according to the manufacturer’s instructions. The final DNA concentration was 1 µg/mL of culture. After 8-24 hours post transfection, sterile filtered valproic acid and 40% tryptone suspended in medium were added to the cultures to final concentrations of 0.5 mM and 0.5% respectively and cultures were then moved to 32°C with continued shaking. Cultures were harvested 5-7 days post-transfection by centrifugation (3,000 × g, 10 min) followed by filtration through 0.2 µm membranes. Antibodies were purified using Protein A affinity chromatography. Briefly, clarified supernatants were loaded onto Protein A columns equilibrated in PBS without calcium and magnesium. Columns were washed with an equilibration buffer and antibodies were eluted with an amine-based elution buffer, pH 2.8, and immediately neutralized with 1 M Tris-HCl, pH 8.5. Eluted fractions were pooled, concentrated, and buffer-exchanged into PBS using dialysis. Protein concentrations were determined by absorbance at 280 nm using calculated extinction coefficients. Antibodies were stored at 4°C for short-term use or at - 80°C in aliquots for long-term storage.

### TNF-α neutralization MTT assay

TNF-α neutralization potency was assessed in a cell-based MTT viability assay. Murine L929 fibroblasts were cultured in DMEM supplemented with 10% fetal bovine serum (FBS), 2 mM L-glutamine, and penicillin–streptomycin at 37°C, 5% CO₂. Cells were seeded into 96-well plates at 1-2 × 10⁴ cells/well and allowed to adhere overnight. Recombinant human TNF-α was prepared at a concentration that produced robust cytotoxicity (typically 0.5-1.0 ng/mL in the presence of actinomycin D). Serial dilutions of each antibody were prepared in assay medium and preincubated with a fixed concentration of TNF-α for 60 min at 37°C. The TNF-α/antibody mixtures were then added to the cells and incubated for 18-24 h. Cell viability was quantified by addition of MTT solution (0.5 mg/mL final concentration) and incubation for 2-4 h, followed by solubilization of formazan crystals in dimethyl sulfoxide (DMSO). Absorbance was measured at 570 nm with a reference wavelength of 630-690 nm. Percent cell protection was calculated relative to wells containing TNF-α alone (0% protection) and wells without TNF-α (100% protection). All measurements were performed in triplicate. Sigmoidal dose-response curves were fitted using a four-parameter logistic model in GraphPad Prism or equivalent software, and half-maximal inhibitory concentrations (IC₅₀) were determined for each antibody. Purified IgG of hInfliximab, infliximab, and adalimumab were included as controls.

### Affinity-capture self-interaction nanoparticle spectroscopy (AC-SINS)

Affinity-capture self-interaction nanoparticle spectroscopy (AC-SINS) was performed to evaluate antibody self-association propensity under low ionic strength conditions. Briefly, gold nanoparticles were coated with polyclonal anti-human IgG capture antibodies and subsequently incubated with test antibodies to generate antibody-conjugated gold nanoparticle complexes. The conjugates were diluted into 20 mM sodium acetate pH 4.3 and incubated at room temperature prior to spectral analysis. Changes in plasmon resonance wavelength Δλmax (nm) were measured using a UV-Vis spectrophotometer, and wavelength shifts were calculated relative to the corresponding buffer control. Increased positive wavelength shifts were interpreted as enhanced self-association and reduced colloidal stability, whereas minimal or negative shifts indicated lower self-interaction propensity and improved developability characteristics. All measurements were performed in triplicate. Purified IgG of hInfliximab, infliximab, and adalimumab were included as controls.

### BVP ELISA

Non-specific binding and polyspecificity were assessed by BVP ELISA. High-binding 96-well plates were coated overnight at 4°C with baculovirus particles diluted in carbonate-bicarbonate buffer (pH 9.6). After washing with PBS pH 7.4, plates were blocked with 0.1% BSA in PBS for 1 hour at room temperature. Test antibodies were diluted in blocking buffer to a concentration of 100 nM and added to the BVP-coated plates in triplicate, followed by incubation for 1 hour at room temperature. After washing, bound human IgG was detected with a horseradish peroxidase-conjugated anti-human Fc-specific secondary antibody and developed with TMB. The reaction was stopped with 0.6 N H₂SO₄, and absorbance was read at 450 nm. For each antibody, a BVP binding ratio was calculated as the ELISA signal at a defined concentration (e.g., 100 nM) normalized to that of secondary antibody background binding. All measurements were performed in triplicate. Higher BVP ratios correspond to increased non-specific binding and have been associated with faster in vivo clearance, whereas low to moderate ratios are considered compatible with favorable pharmacokinetics. Purified IgG of hInfliximab, infliximab, and adalimumab were included as controls.

### Polyspecificity evaluation of purified IgGs of targeted amino-acid mutant clones of hInBG4 by HSP90-hFc and dsDNA ELISA

Microtiter wells were coated with HSP90-hFc (10 µg/mL) or dsDNA (2 µg/mL) in PBS at 4°C overnight. Following washing and blocking, purified IgGs (100 nM) were added to the wells for incubation at room temperature for 1 hour. After washing, bound IgG was detected with peroxidase conjugated goat anti-human IgG Fcγ specific antibody (Jackson ImmunoResearch). All measurements were performed in triplicate. The ratio was calculated by dividing the signal of each clone by the signal of blank control (0.1% BSA/PBS). Purified IgGs of hInfliximab and adalimumab were also included as controls in this study.

### Polyspecificity evaluation of purified IgGs of targeted amino-acid mutant clones of hInBG4 by flow cytometry

For polyspecificity evaluation by flow cytometry, streptavidin beads coated with biotinylated HSP90-hFc were incubated with purified IgGs 100 nM. Beads were washed with FACS and bound IgG was detected with AlexaFluor 647-conjugated goat anti-human IgG Fcγ specific antibody (Jackson ImmunoResearch). All measurements were performed in triplicate. The ratio was calculated by dividing the signal of each clone by the signal of blank control (FACS only). Purified IgGs of hInfliximab and adalimumab were also included as controls in this study.

### SE-HPLC

Size variants and aggregation were analyzed by SE-HPLC. Antibodies were diluted to 0.5 mg/mL in mobile phase (e.g., 50 mM sodium phosphate, 400 mM NaCl, pH 6.8). Samples (typically 10 µg per injection) were injected onto a TSKgel UP-SW3000size-exclusion column equilibrated in the same buffer. Chromatography was performed at room temperature with a flow rate of 0.35 mL/min, and elution was monitored by UV absorbance at 215 and 280 nm. Samples were bracketed using a NIST monoclonal antibody (NISTmAb) reference material (RM 8671) run at regular intervals to monitor system performance and provide a reference for monomer peak position. The percentage of monomer and high-molecular-weight species (HMWS) was determined by integration of the chromatogram using ChemStation. Symmetrical monomer peaks with minimal tailing and low HMWS content were interpreted as evidence of favorable chromatographic and colloidal behavior.

### Expression quantitation

Relative expression levels in transient HEK293 cultures were quantified using a biolayer interferometry (BLI) IgG capture assay on an Octet instrument (Sartorius). Clarified culture supernatants were harvested as described above and diluted in kinetics buffer (PBS with 0.02% Tween-20 and 0.1% BSA). Protein A biosensors (Sartorius 18-5010) were pre-equilibrated in a kinetics buffer and then loaded with samples in a parallel format. A standard curve was generated using purified human IgG1 at known concentrations spanning the expected range of expression. The maximum binding response (nm shift) for each sample was interpolated against the standard curve to estimate antibody titer in the culture supernatant. Expression levels for engineered clones were compared to those of hInfliximab, which served as a parental humanized control.

### Octet kinetics at pH 7.4 and 5.8

TNF-α binding kinetics were measured by BLI using an Octet instrument at two pH conditions. Recombinant human TNF-α was biotinylated using NHS-LC-biotin according to the manufacturer’s instructions and desalted, then diluted into a kinetics buffer. Streptavidin (SA) biosensors were pre-equilibrated in kinetics buffer at pH 7.4 and then loaded using a solution containing 5 nM biotinylated TNF-α for 300 seconds. Kinetic assays were performed at 30°C in 96-well plates containing antibody solutions at multiple concentrations in the kinetics buffer. Association was monitored by dipping TNF-α-loaded sensors into wells containing serial dilutions of each antibody for a fixed time (120 s). Dissociation was then monitored by transferring sensors into wells containing kinetics buffer at either pH 7.4 or pH 5.8, prepared by adjusting buffer components while maintaining ionic strength. Sensorgrams were reference-subtracted using buffer-only controls and analyzed with the Octet data analysis software. Global fitting of association and dissociation phases across multiple concentrations was performed using a 1:1 Langmuir binding model to obtain association rate constants (ka), dissociation rate constants (kdis), and equilibrium dissociation constants (KD = kdis/ka). For clones with extremely slow off-rates at pH 7.4, kdis values below the reliable limit of detection were reported as <1.0 × 10⁻⁷ s⁻¹. The ratio of kdis at pH 5.8 to that at pH 7.4 was used as a quantitative measure of pH-dependent antigen dissociation.

### In silico assessment of T-cell epitope content

Predicted T-cell epitope content was evaluated in silico using the IEDB Deimmunization tool (https://tools.iedb.org/deimmunization/). Variable heavy and light chain sequences of chimeric infliximab, hInfliximab, hInBG4 and the engineered variants LW2Y, LW2YR2S, HF1Y and LW2YHR1K were analyzed using default settings. For each sequence, immunogenicity scores were calculated based on predicted Major Histocompatibility Complex (MHC) class II binding across the reference Human Leukocyte Antigen panel, and the overall scores and distribution of predicted high-risk epitopes were compared qualitatively between the parental and engineered antibodies.

## RESULTS

### Humanization and STEM™-mediated engineering of infliximab

The overall engineering strategy and library design are summarized in Figure 1. We first humanized the variable regions of infliximab by grafting the CDRs onto selected human germline frameworks while preserving key canonical residues. The resulting humanized infliximab (hInfliximab) retained TNF-α binding but showed reduced potency in a TNF-α neutralization assay compared with the parental chimeric antibody. To recover and improve functional activity, we applied our STage-Enhanced Maturation (STEM™) workflow to hInfliximab and obtained an affinity-matured variant, hInBG4, that exhibited enhanced cell protection in a TNF-α-induced cytotoxicity (MTT) assay with low non-specific binding by BVP ELISA (Table 1S, Figure 2A–2B). However, hInBG4 still showed strong self-association like the original infliximab in AC-SINS (Figure 2C).

**Figure 1.**
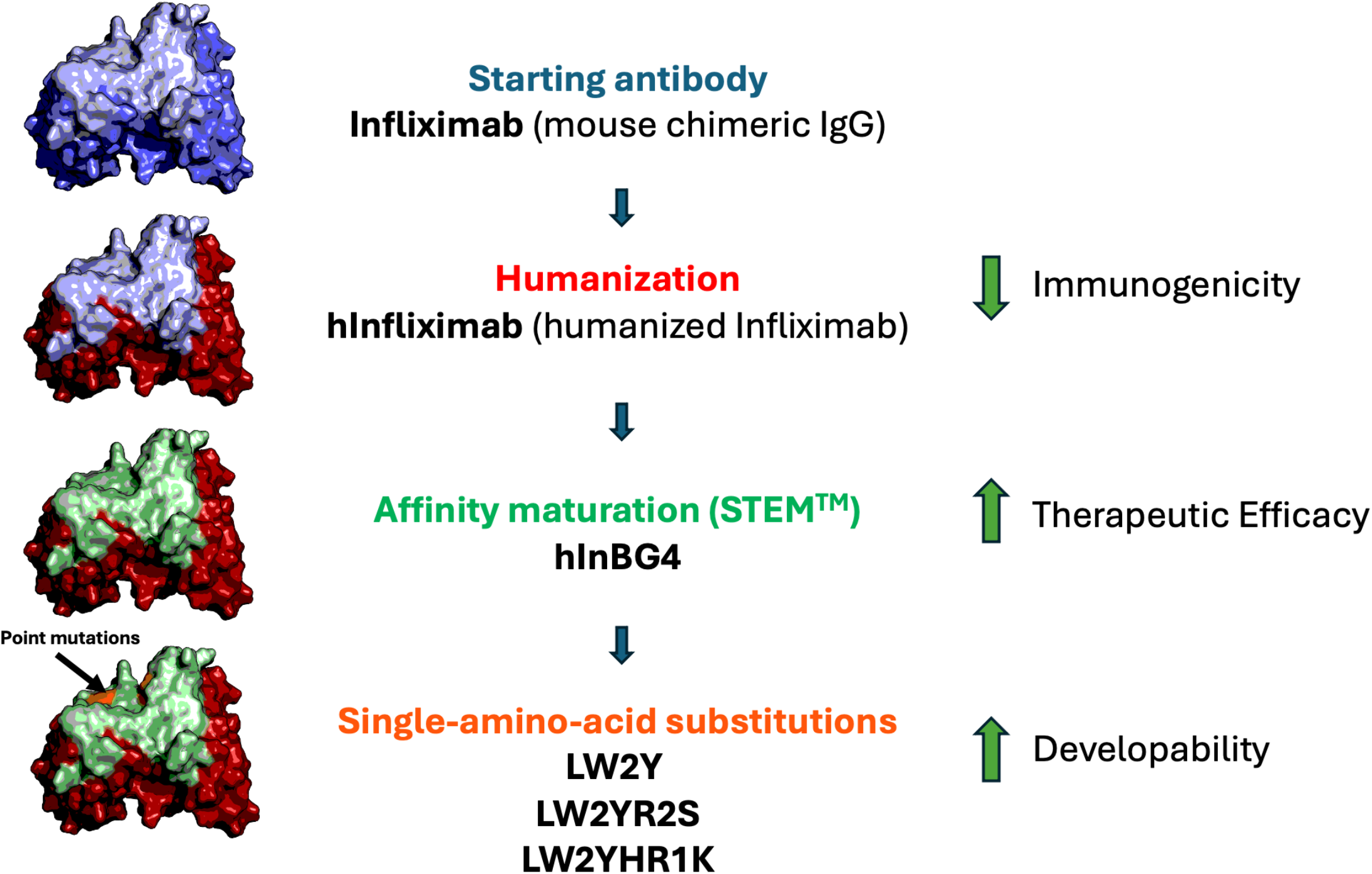
Engineering workflow for infliximab-derived variants. Infliximab was humanized, affinity matured using STEM™ technology, and subsequently refined by targeted-amino-acid substitutions to generate clones with improved potency and developability. Murine CDRs and frameworks are shown in light blue and blue, respectively. Human frameworks are shown in red, and engineered CDRs are shown in green.

**Figure 2.**
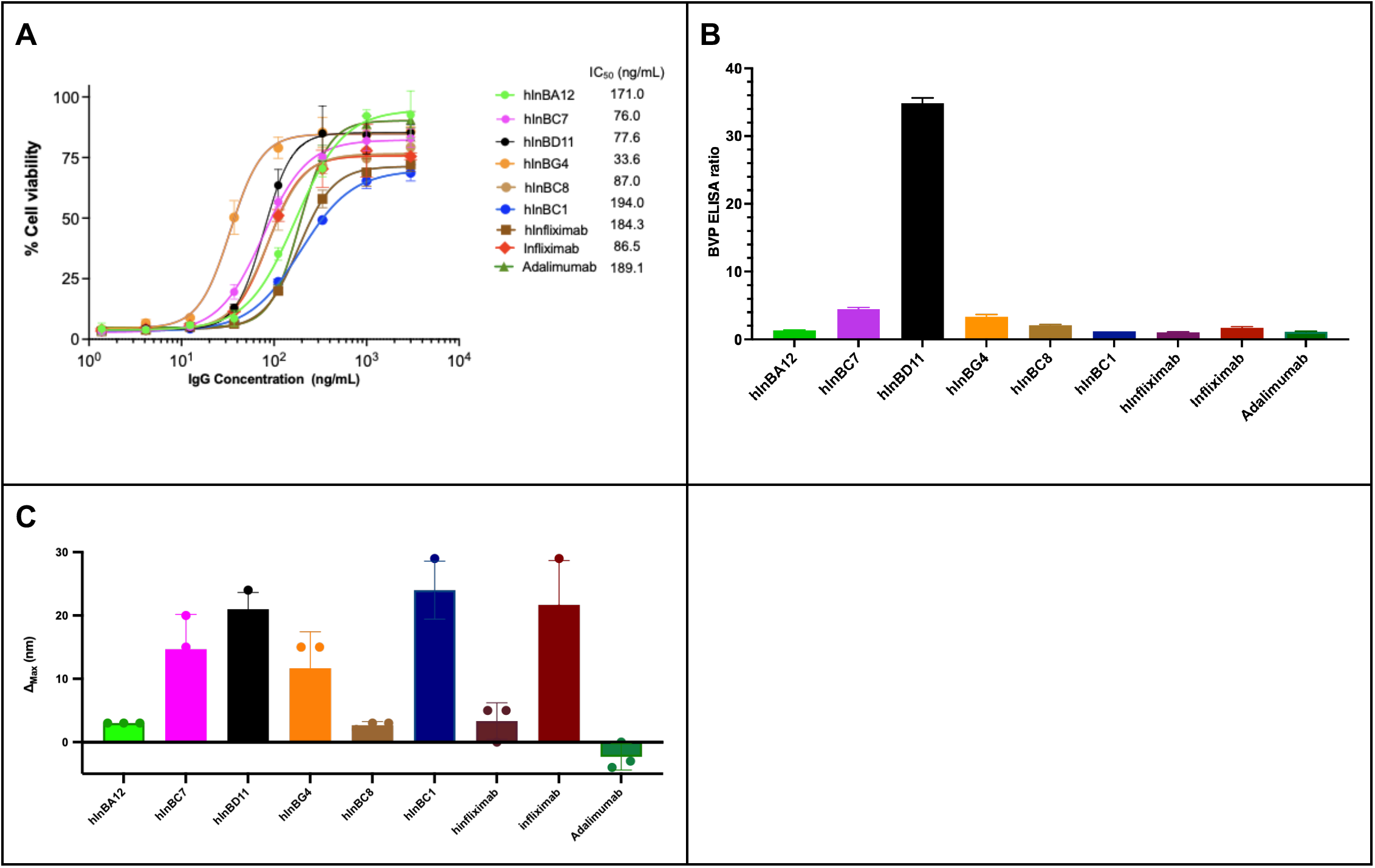
STEM™-selected humanized infliximab variants show improved neutralization but persistent self-association. (A) MTT assay of purified IgG clones selected by STEM™ engineering. Clone hInBG4 showed the lowest IC₅₀value among the tested STEM™-selected clones, with hInfliximab, infliximab, and adalimumab included as controls. (B) Polyspecificity of purified IgG clones selected by STEM™ engineering was assessed by BVP ELISA. All clones except hInBD11 showed low BVP ELISA ratios. (C) Purified IgGs were assessed by AC-SINS. Selected clones showed relatively high Δλmax values, similar to the original infliximab and higher than adalimumab. For panels A–C, data are shown as mean ± SD of triplicate measurements.

### Orthogonal polyspecificity assays of crude Fabs

Polyspecificity of engineered Fabs was also tested by HSP90 ELISA, dsDNA ELISA, and HSP90-coated bead flow cytometry assays. HSP90 ELISA and dsDNA ELISA showed high correlation (Figure 1S) while HSP90-coated bead flow cytometry assay did not show good correlation with these ELISA results (Figure 2S).

### Engineered variants show improved TNF-α neutralization potency

Using hInBG4 as the parental scaffold, we then constructed a panel of targeted amino-acid CDR mutants. TNF-α-induced cytotoxicity was measured in the presence of serial dilutions of each antibody (Figure 3A). Subsequent developability profiling by AC-SINS, BVP ELISA, SE-HPLC (Figure 3B-3E) and transient expression led to the identification of three clones - LW2Y, LW2YR2S, and LW2YHR1K - for detailed characterization. Compared with hInfliximab (IC₅₀ 166.1 ng/mL) and chimeric infliximab (96.8 ng/mL), all three engineered variants displayed substantially lower IC₅₀ values, ranging from 33.2 to 45.4 ng/mL (Figure 3A, Table 1). Specifically, LW2YHR1K showed the greatest potency (33.2 ng/mL), followed by LW2Y (34.5 ng/mL), and LW2YR2S (45.4 ng/mL). Adalimumab, included as a benchmark fully human anti-TNF-α antibody, exhibited an IC₅₀ of 167.0 ng/mL, comparable to hInfliximab and higher than all three engineered clones.

**Figure 3.**
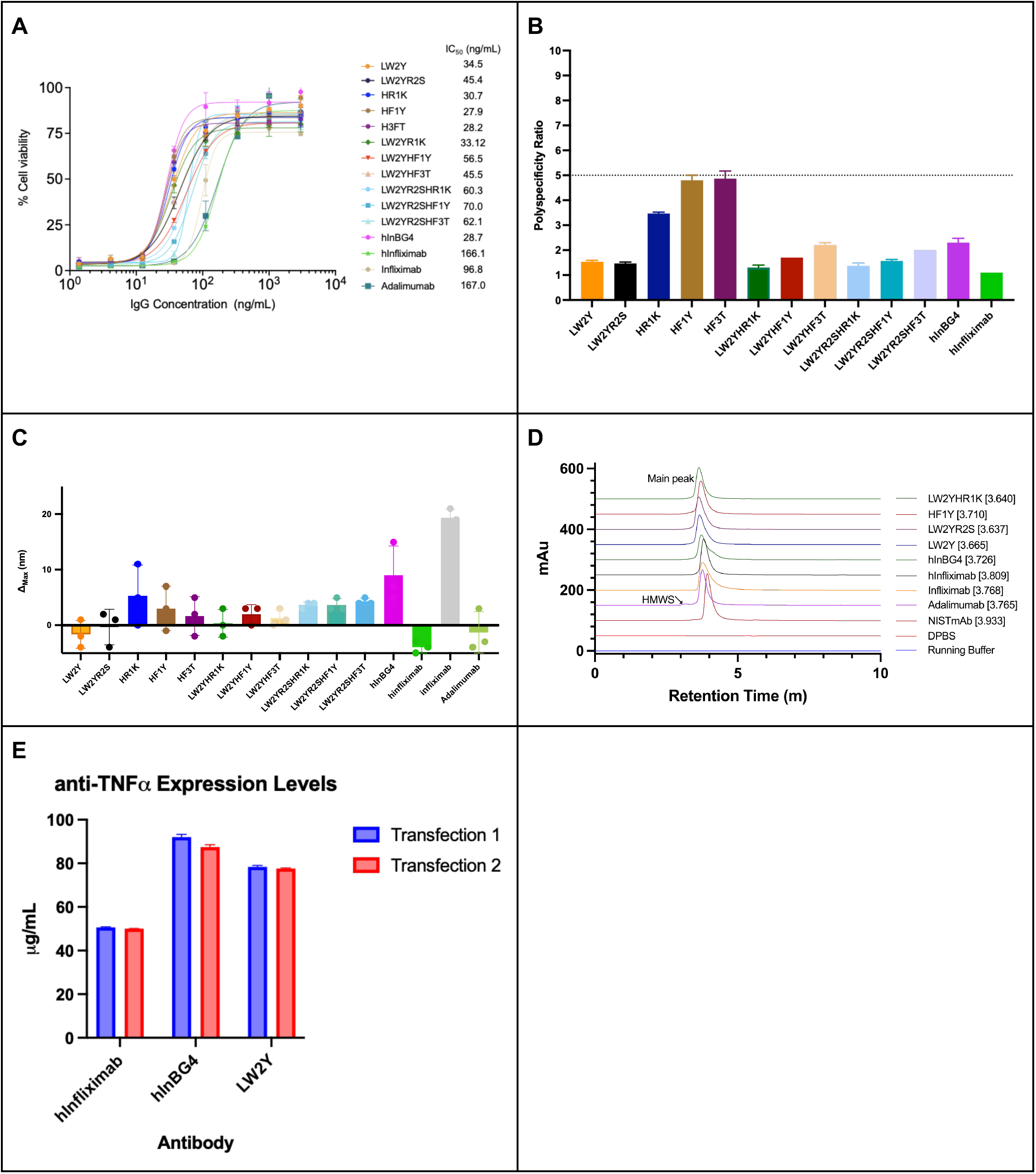
Targeted amino-acid CDR substitutions improve potency and developability of hInBG4-derived variants. (A) MTT assay of hInBG4 variants with targeted amino-acid substitutions. hInBG4 and its variants showed increased neutralization potency compared with hInfliximab, infliximab, and adalimumab. (B) BVP ELISA analysis of hInBG4 variants with targeted amino-acid substitutions. BVP ELISA ratios of LW2Y, LW2YR2S, LW2YHR1K, LW2YHF1Y, LW2YR2SHR1K, and LW2YR2SHF1Y were less than 2.0. (C) AC-SINS analysis of hInBG4 variants with targeted amino-acid substitutions. LW2Y, LW2YR2S, and LW2YHR1K showed low Δλmax values comparable to that of adalimumab. (D) SE-HPLC profiles of infliximab-derived antibodies. Chromatograms are shown as vertically offset traces for clarity. The monomer main peak and HMWS region are indicated. Infliximab and hInBG4 exhibited peak tailing, whereas engineered variants showed sharper, more symmetrical monomer peaks. Main-peak retention times are shown in brackets for each sample. Adalimumab and NISTmAb RM 8671 were included as comparators; DPBS and running buffer served as blanks. (E) Relative expression levels of engineered clones compared with parental humanized infliximab. Selected antibodies were transfected in duplicate and expression was quantified by Octet IgG quantification. Each sample was measured in duplicate. Median and range are shown. Engineered clones showed higher expression levels than hInfliximab. For panels A–C, data are shown as mean ± SD of triplicate measurements.

**Table 1.** Summary of potency and developability metrics for infliximab-derived antibodies.

| | MTT IC<sub>50</sub> (ng/mL) | AC-SINS $\Delta\lambda_{\max}$ (nm) | BVP ELISA ratio |
| --- | --- | --- | --- |
| LW2Y | 34.5 | -1.67 | 1.5 |
| LW2YR2S | 45.4 | -0.33 | 1.5 |
| HR1K | 30.7 | 5.33 | 3.5 |
| HF1Y | 27.9 | 3.00 | 4.8 |
| HF3T | 28.2 | 1.67 | 4.9 |
| LW2YHR1K | 33.2 | 0.33 | 1.3 |
| hInBG4 | 42.0 | 9.00 | 2.3 |
| hInfliximab | 166.1 | -4.00 | 1.1 |
| Infliximab | 96.8 | 19.33 | 1.3 |
| Adalimumab<sup>#</sup> | 167.0 | -1.33 | 1.6 |

<sup>#</sup> in-house expressed infliximab and adalimumab were included as controls

### Improved self-association and non-specific binding profiles of targeted amino-acid CDR mutants of hInBG4 assessed by AC-SINS and BVP ELISA

BVP ELISA ratios for these engineered variants remained within the range of 1.3-4.9 (Figure 3B and Table 1). LW2Y (1.5), LW2YR2S (1.5), LW2YHR1K (1.3) were comparable to infliximab (1.3) and adalimumab (1.6), whereas HR1K, HF1Y and HF3T showed higher values (3.5 to 4.9). Orthogonal polyspecificity assays with HSP90 ELISA, dsDNA ELISA, HSP90-coated bead flow cytometry results correlated very well with the results of BVP ELISA (Figure 3S and 4S).

Developability profiling by AC-SINS revealed that chimeric infliximab and an engineered humanized infliximab clone hInBG4 exhibited relatively high plasmon peak shifts (Δλmax values of 19.33 and 9.00 nm), respectively, consistent with an increased propensity for self-association (Figure 3C and Table 1). In contrast, the engineered clones LW2Y, LW2YR2S, and LW2YHR1K showed reduced AC-SINS responses, with Δλmax values of -1.67 - 0.33 nm (Figure 3C and Table 1). LW2Y and LW2YR2S displayed the lowest values (Δλmax value -1.67 and -0.33 nm) similar to adalimumab (Δλmax value -1.33 nm).

### SE-HPLC and transient expression support improved developability

SE-HPLC analysis demonstrated that chimeric infliximab and hInBG4 exhibited main peaks accompanied by shoulder and tailing peaks (Figure 3D). In contrast, LW2Y, LW2YR2S, and LW2YHR1K showed sharp, symmetrical main peaks with minimal tailing. In-house produced adalimumab exhibited more HMWS than the other antibodies analyzed. To assess expression, we transiently transfected HEK293 cells with each antibody and quantified secreted IgG by Octet IgG capture. Engineered clones expressed at higher levels than hInfliximab, with LW2Y showing the largest increases (Figure 3E).

### pH-dependent TNF-α binding identifies LW2Y as a pH-sensitive lead

To explore pH-dependent antigen binding, we measured TNF-α kinetics by BLI at pH 7.4 and 5.8, using identical association conditions followed by dissociation in buffers of either pH (Figure 4, Table 2). At pH 7.4, LW2Y exhibited extremely high affinity, with an apparent KD < 1 pM and an off-rate below 1.0 × 10⁻⁷ s⁻¹. At pH 5.8, LW2Y displayed a markedly increased off-rate (1.8 × 10⁻⁴ s⁻¹), corresponding to a >1800-fold acceleration relative to the assay-detection-limited off-rate at pH 7.4. LW2YR2S also showed increased off-rates at pH 5.8, albeit to a lesser extent, while hInfliximab and infliximab exhibited more modest pH-dependent changes. Adalimumab demonstrated very high affinity at pH 7.4 (KD 0.012 nM) and a substantial increase in off-rate at pH 5.8 (Table 2). The combination of ultra-slow dissociation at pH 7.4 and pronounced acceleration at pH 5.8 distinguished LW2Y from both the parental and comparator antibodies.

**Figure 4.**
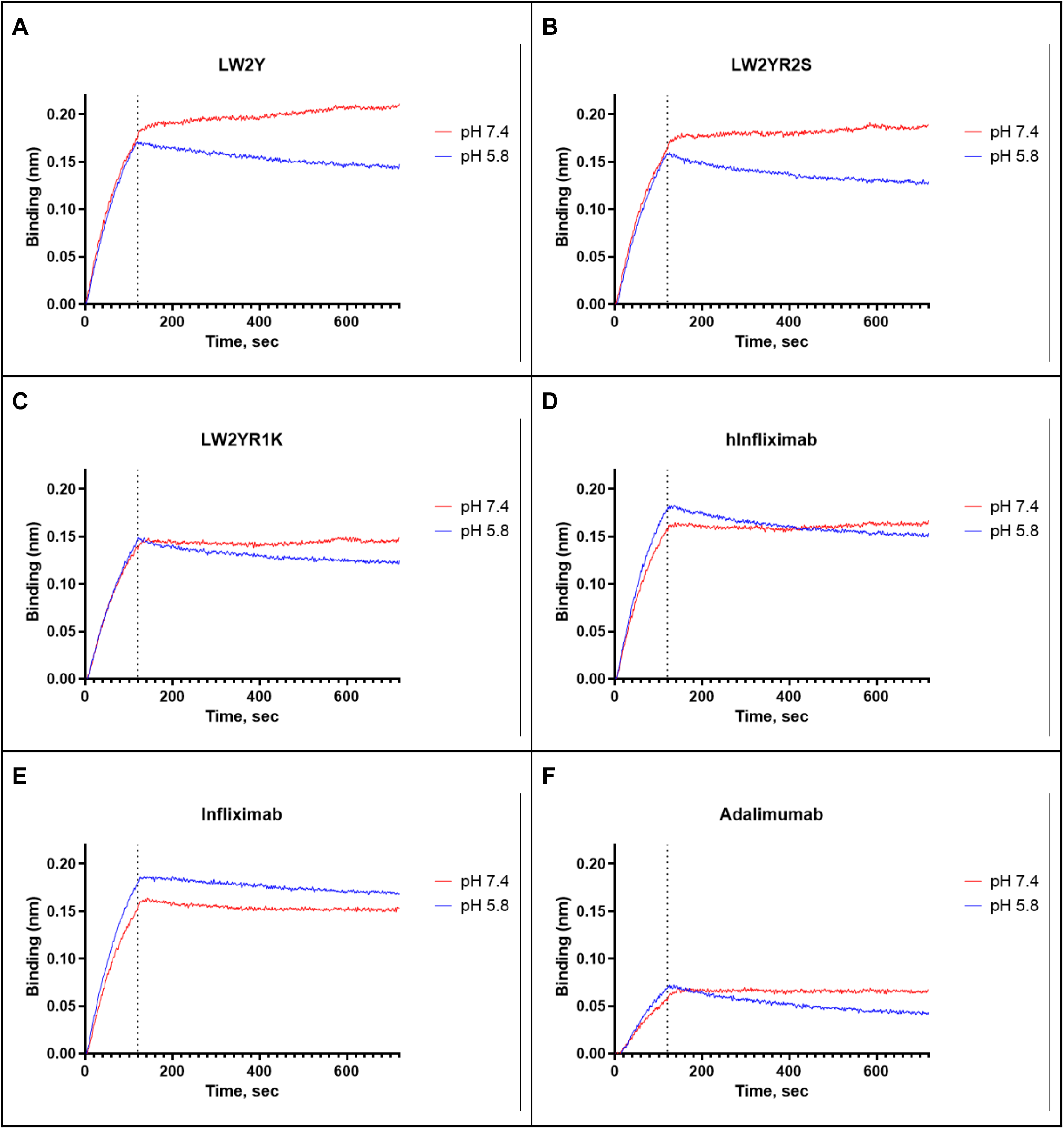
pH-dependent TNF-α dissociation measured by BLI. TNF-α was captured on streptavidin biosensors and antibodies were associated at pH 7.4. Dissociation was monitored in the buffer at pH 7.4 or pH 5.8. Traces are shown for the indicated antibodies associated at a concentration of 5 nM.

**Table 2.** TNF-α binding kinetics at pH 7.4 with dissociation measured at pH 7.4 or pH 5.8 by BLI.

| Clone | K<sub>D</sub> (nM) pH 7.4 | k<sub>a</sub> (M<sup>-1</sup> s<sup>-1</sup>) pH 7.4 | k<sub>dis</sub> (s<sup>-1</sup>) pH 7.4 | k<sub>dis</sub> (s<sup>-1</sup>) pH 5.8 | k<sub>dis</sub> pH 5.8/k<sub>dis</sub> pH 7.4<sup>#</sup> |
| --- | --- | --- | --- | --- | --- |
| LW2YHR1K | 0.539 | 1.76E+05 | 9.49E-05 | 1.77E-04 | 1.87 |
| LW2YR2S | 0.056 | 1.74E+05 | 9.82E-06 | 1.80E-04 | 18.33 |
| LW2Y | <0.001 | 1.06E+05 | <1.0E-07 | 1.80E-04 | >1800 |
| hInfliximab | 0.752 | 2.12E+05 | 1.60E-04 | 2.82E-04 | 1.76 |
| Infliximab | 1.009 | 2.17E+05 | 2.19E-04 | 1.51E-04 | 0.69 |
| Adalimumab | 0.012 | 1.72E+05 | 2.01E-06 | 4.35E-04 | 216.42 |

<sup>#</sup>The k<sub>dis</sub> pH 5.8/k<sub>dis</sub> pH 7.4 ratio reports the fold-change in dissociation rate at acidic vs neutral pH; larger values indicate faster dissociation at pH 5.8 (greater pH-dependent antigen release). For LW2Y, k<sub>dis</sub> at pH 7.4 was below the assay's reliable detection limit and is reported as <1.0 × 10<sup>-7</sup> s<sup>-1</sup>; therefore, the ratio is shown as a lower bound (>1800).

### In silico immunogenicity assessment of engineered variants

To examine potential differences in T-cell epitope content, we analyzed the VH and VL sequences of infliximab, hInfliximab and the STEM™-engineered variants (hInBG4, LW2Y, LW2YR2S, and LW2YHR1K) using the IEDB Deimmunization tool (peptide mutant prediction mode) with a median percentile rank threshold of 20. For the light chain of chimeric infliximab, a single peptide (DILLTQSPAILSVSP) was identified with a median percentile rank of 16.5, indicating a predicted MHC class II–restricted epitope at the chosen threshold. In contrast, no peptides meeting this threshold were detected in the light chains of hInfliximab, hInBG4, LW2Y, LW2YR2S or LW2YHR1K, nor in any of the corresponding heavy-chain sequences. Within the limits of this in silico approach, these data suggest that humanization and CDR-level engineering did not introduce additional high-risk predicted T-cell epitopes and removed one predicted light-chain epitope present in infliximab.

## DISCUSSION

In this study, the chimeric anti-TNF-α antibody infliximab was humanized and subsequently optimized through focused CDR engineering using the STEM™ platform and targeted amino-acid substitutions to generate variants with improved potency and developability (Figure 1). Starting from humanized infliximab (hInfliximab), we obtained an affinity-matured intermediate (hInBG4) that exhibited greater inhibition of TNF-α-induced cellular cytotoxicity than infliximab and adalimumab (Figure 2A) and clean BVP ELISA ratio (Figure 2B) but with modestly elevated AC-SINS value (Figure 2C). Then, we identified three engineered clones of hInBG4 - LW2Y, LW2YR2S, and LW2YHR1K - that showed consistently lower IC₅₀ values in a TNF-α-induced cytotoxicity assay than both chimeric infliximab and adalimumab (Figure 3A) with favorable biophysical profiles, including acceptable BVP ELISA behavior (Figure 3B), reduced self-association by AC-SINS (Figure 3C), cleaner SE-HPLC chromatograms (Figure 3D), and higher transient expression (Figure 3E). Collectively, these data support the conclusion that STEM™-guided humanization and fine-scale CDR optimization can convert a clinically validated but immunogenic chimeric antibody into a panel of humanized biobetter candidates with improved functional activity and developability.

The humanization strategy was designed to reduce potential immunogenicity risk associated with the murine variable regions of infliximab while maintaining TNF-α binding. Anti-drug antibodies against infliximab variable domains are associated with secondary loss of response and infusion reactions in patients [1, 2]. The humanized scaffold described here preserves overall paratope architecture while replacing murine frameworks with human germline sequences, an approach that has historically reduced immunogenicity for other therapeutics [1]. At the same time, hInfliximab showed reduced neutralization potency relative to the chimeric parent (Figure 2A), highlighting a common practical challenge in humanization campaigns and underscoring the need for subsequent affinity and functional maturation. By stepwise affinity maturation through STEM™ library design and selection, followed by direct functional screening in a cell-based TNF-α neutralization assay, we not only recovered but surpassed the neutralizing activity of infliximab and adalimumab while maintaining a humanized framework expected to be more compatible with chronic administration. Consistent with this design goal, an in silico analysis using the IEDB Deimmunization tool suggested that humanization removed a predicted light-chain MHC class II epitope present in infliximab and that subsequent CDR-level engineering did not introduce new high-risk clusters in the variable regions, within the limits of this computational approach.

A key question in advancing any lead antibody is whether its biophysical properties fall within ranges that have proven compatible with clinical development. Large-scale analyses of marketed and clinical-stage antibodies define an approximate “developability envelope” for self-association, non-specific binding, and colloidal stability [4–6]. AC-SINS, in particular, has emerged as a sensitive, high-throughput readout of weak antibody–antibody interactions that correlate with aggregation liability, high viscosity at formulation concentrations, and other unfavorable colloidal behaviors [7–10]. In this context, the double-digit AC-SINS shift observed for infliximab is consistent with increased self-association propensity, whereas the low single-digit shifts for LW2Y, LW2YR2S, and LW2YHR1K place these engineered variants closer to the favorable region observed for many clinical-stage molecules (Figure 3C and Table 1). Importantly, this developability dimension is particularly relevant for infliximab because reversible self-association has been mechanistically linked to Fab-mediated interfaces observed structurally and supported by solution behavior [3]. Thus, the improvements in AC-SINS and SEC profiles seen in the optimized variants are consistent with the idea that targeted CDR substitutions can modulate the self-interaction landscape of an infliximab-derived antibody while preserving antigen specificity.

Non-specific binding, as measured by BVP ELISA, is another important parameter with pharmacokinetic implications. Hötzel and colleagues reported that higher baculovirus particle binding correlates with faster, non-target-mediated clearance in cynomolgus monkeys and proposed BVP ELISA as an early risk-mitigation tool [11]. Subsequent work has reinforced the association between polyspecificity/polyreactivity and poor pharmacokinetics or increased off-target interactions [10, 12–15, 26]. In the present panel, infliximab and adalimumab exhibited low BVP ratios (1.3 and 1.6, respectively), whereas hInBG4 showed slightly elevated binding (2.3) (Table 1). Overall, five out of six engineered clones of hInfliximab tested and nine out of 11 clones of targeted amino-acid mutants of hInBG4 showed acceptable BVP ratios (<5.0) suggesting that the STEM™ workflow with negative selection with PSRs paralleled potency gains and avoidance of excessive polyreactivity. Although in vivo PK studies will ultimately be required, the combination of low AC-SINS and acceptable BVP binding supports the view that LW2Y and LW2YR2S, in particular, occupy an acceptable developability space for further preclinical development.

Orthogonal specificity assays of purified IgGs using HSP90 ELISA, dsDNA ELISA and HSP90-coated bead flow results correlated well with the results of BVP ELISA although the HSP90-coated bead flow results of crude Fabs did not show good correlation (Figures 1S-4S). Crude Fabs showed generally lower signal ratios (<3.0) except for adalimumab and another clone on the bead platform. This likely results from stronger avidity dependence in the bead assay and interference from bacterial proteins in crude samples.

SE-HPLC provided a complementary readout of aggregation and size variants. High monomer content and minimal high-molecular-weight species are fundamental requirements for clinical development, and SE-HPLC is routinely used as part of broader developability assessments [4–6]. Infliximab and hInBG4 displayed shoulders and tailing consistent with on-column interactions and/or size heterogeneity, which may align with their higher self-association propensity. In contrast, the engineered variants showed sharper, more symmetrical monomer peaks with reduced tailing (Figure 3D). Heat treatment during STEM™-engineering also selects clones with more efficient folding that tend to express better than its parent antibody (Figure 3E). Together with increased expression yields in transient HEK293 systems, these findings support improved manufacturability and motivate future formulation studies, including assessment of high-concentration behavior relevant to potential subcutaneous administration.

A distinguishing feature of this optimized panel is the pH dependence of TNF-α binding, most notably for LW2Y (Figure 4A and 4B, Table 2). Antibodies engineered to bind antigen tightly at neutral pH but release it under mildly acidic endosomal conditions have been proposed to enhance antigen clearance and mitigate target-mediated drug disposition by leveraging FcRn-mediated recycling of IgG [16, 27]. In this conceptual framework, antigen-antibody complexes internalize via fluid-phase or target-mediated uptake; the antibody dissociates from antigen in the endosome, allowing antigen to be degraded while IgG is salvaged by FcRn and returned to circulation [16, 27]. For LW2Y, the combination of an extremely slow off-rate at pH 7.4 with a markedly accelerated dissociation at pH 5.8 is consistent with a recycling-like antigen release profile. This behavior provides a clear point of comparison with prior anti-TNF-α pH-switch engineering efforts reported for adalimumab/Humira-derived antibodies, including a monovalent adalimumab variant designed for endosomal TNF-α release and reported to reduce immunogenicity in mice [17], as well as histidine-based engineering and in vitro/computational studies of pH-dependent TNF-α dissociation [18]. In that context, LW2Y is notable in that its strong pH-dependent dissociation was achieved through CDR-level engineering on an infliximab-derived IgG while simultaneously improving self-association and manufacturability metrics.

The extent to which LW2Y’s pH profile translates into superior pharmacokinetics, antigen clearance, or improved durability of neutralization will depend on the interplay between antigen binding kinetics, FcRn engagement, and TNF-α turnover. FcRn mediates IgG recycling by binding at acidic pH and releasing IgG at neutral pH, thereby extending half-life [27]. Modeling and in vivo studies indicate that non-specific binding, target-mediated clearance, and FcRn interactions collectively shape the pharmacokinetics of humanized antibodies [14, 26]. In the present work, LW2Y’s pH-dependent TNF-α binding was achieved without Fc modifications, suggesting at least two practical next steps: (i) direct in vivo evaluation of pharmacokinetics and TNF-α clearance to test whether CDR-encoded pH sensitivity confers measurable advantages over infliximab and adalimumab; and (ii) exploration of combinations with Fc variants that enhance FcRn binding, which could further amplify recycling behavior and exposure while maintaining favorable developability.

Despite these encouraging results, several limitations should be acknowledged. First, developability assessment focused on a subset of in vitro assays (AC-SINS, BVP ELISA, SE-HPLC, and transient expression) and did not include viscosity measurements at formulation-relevant concentrations, long-term stability studies, or additional orthogonal polyspecificity panels that can further refine developability risk [5–6, 10, 12–13, 15]. Second, immunogenicity risk was addressed through humanization and in silico T-cell epitope prediction rather than experimental T-cell assays or in vivo immunogenicity studies; therefore, the absence of elevated predicted scores should be considered supportive but not definitive. Third, all functional evaluations were performed in vitro; consequently, whether the combined potency, developability improvements, and pH-dependent binding translate into superior efficacy, safety, and pharmacokinetics in vivo remains to be established.

In summary, this work demonstrates that humanization, STEM™-mediated stepwise affinity maturation, and fine CDR optimization can yield infliximab-derived biobetters that combine increased TNF-α neutralization potency with developability metrics aligned with clinical-stage benchmarks for self-association, non-specific binding, and aggregation [4–13, 15]. Among the engineered variants, LW2Y is particularly notable for its combination of high affinity at physiological pH, pronounced pH-dependent antigen release, low AC-SINS signal, acceptable BVP binding, favorable SE-HPLC behavior, and improved expression. These attributes motivate further preclinical evaluation of LW2Y as a potential next-generation anti-TNF-α therapeutic and support the broader premise that coupling CDR-encoded pH sensitivity with developability-guided engineering may provide a viable route to more durable and convenient TNF-α inhibitors [3, 14, 16–18, 26, 27].

## ACKNOWLEDGMENTS

The authors used ChatGPT for language editing and manuscript review. The authors reviewed and approved all content and take full responsibility for the final manuscript.

## DATA AVAILABILITY

The datasets used and/or analyzed during the current study are available from the corresponding author on reasonable request.

## AUTHORS’ CONTRIBUTIONS

P.D.E. conducted panning of CDR mutant libraries and screening of isolated clones, BVP ELISA and other polyspecificity assays to assess polyreactivity, and affinity measurement by Octet RED 96e and analyzed Ka and Kdis at pH 7.4 and pH 5.8. K.C.E. designed and constructed CDR mutant libraries, and panning protocols to support the work of P.D.E.. J.K.F. procured all antibodies and handled expression, purification, SE-HPLC, methods development, and figures. A.S. performed manufacturability and biophysical risk assessment by AC-SINS. L.Y.E. and Y. H. performed cloning work necessary for the conversion of Fabs to IgG. S.C.J.O. provided strategic oversight for the project, guided experimental priorities, coordinated cross-team collaboration, and contributed to data interpretation and manuscript refinement. T.M. initiated and managed the whole project, handled humanization of infliximab, single amino acid mutant design and cloning, and wrote the manuscript, with all authors providing comments and edits.

## ETHICS AND CONSENT STATEMENT

Not applicable.

## ANIMAL RESEARCH STATEMENT

Not applicable.

## Figure legends

**Figure 1S.**
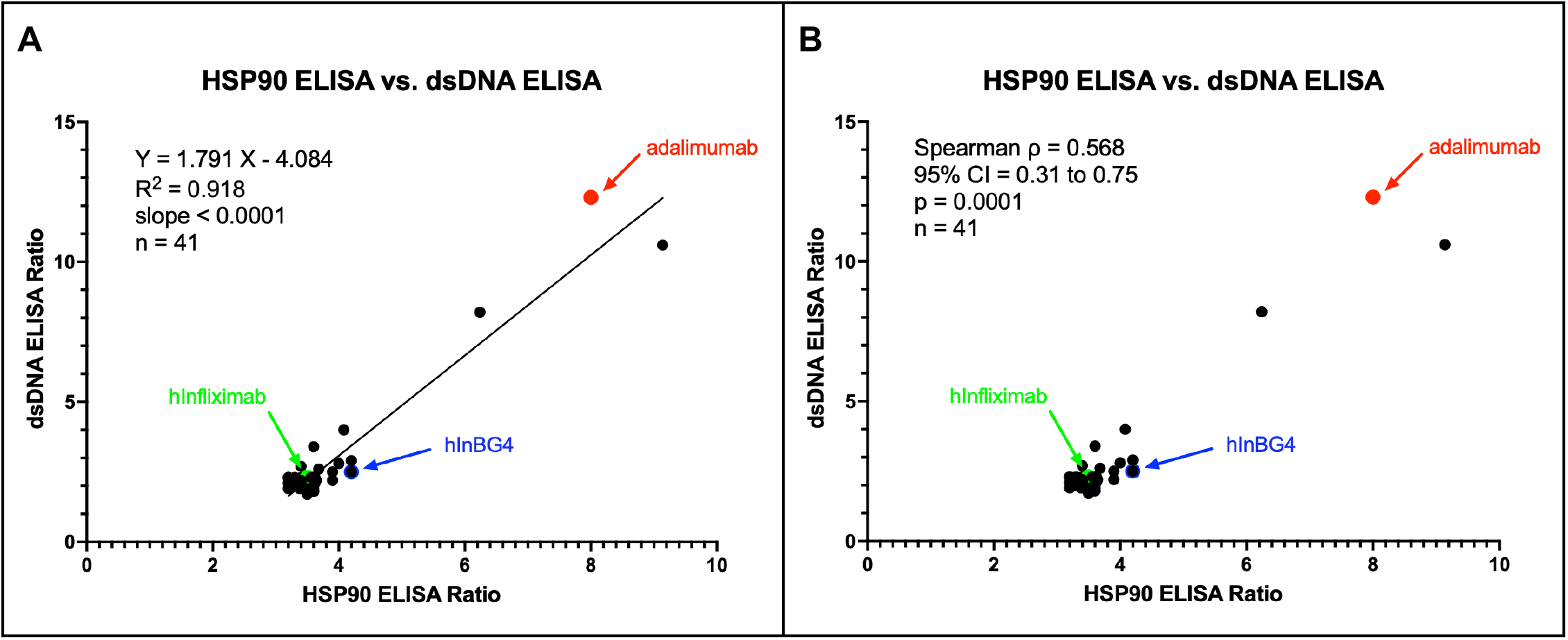
Orthogonal polyspecificity assays of crude Fabs selected by STEM™ engineering (crude Fabs of hInfliximab and adalimumab were included as controls): Polyspecificity of clones selected by STEM™ engineering was tested using HSP90 ELISA and dsDNA ELISA. There was a strong correlation between the HSP90 ELISA and dsDNA ELISA ratios for the crude Fab.

**Figure 2S.**
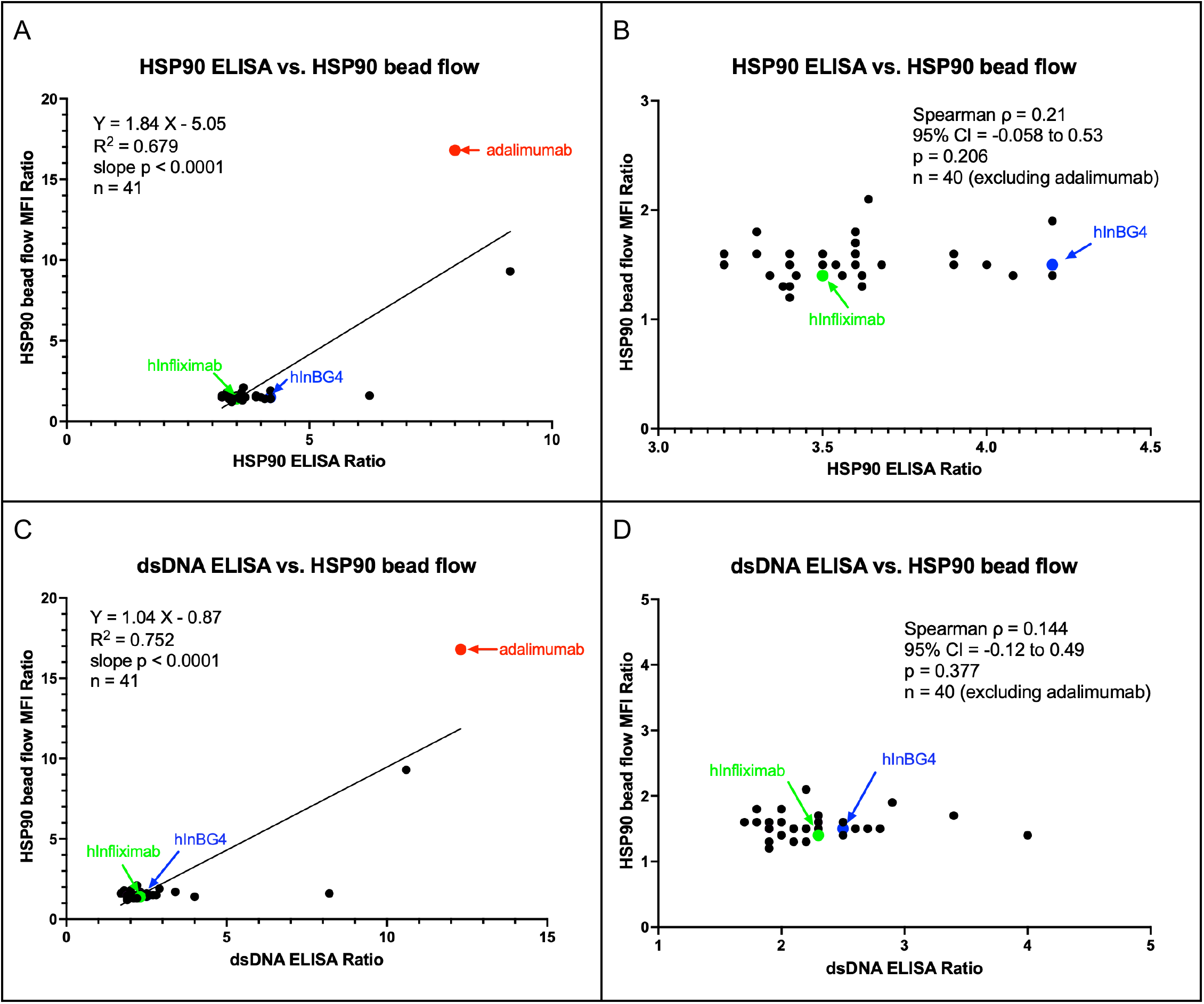
Orthogonal polyspecificity assays of crude Fabs selected by STEM™ engineering (crude Fabs of hInfliximab and adalimumab were included as controls): Polyspecificity of clones selected by STEM™ engineering was tested using biotinylated HSP90-hFc captured to streptavidin-beads in flow cytometry (HSP90 bead flow) and compared with the results of HSP90 ELISA and dsDNA ELISA. HSP90 bead flow ratios were generally low and showed no correlation with HSP90 ELISA, dsDNA ELISA for the crude Fabs.

**Figure 3S.**
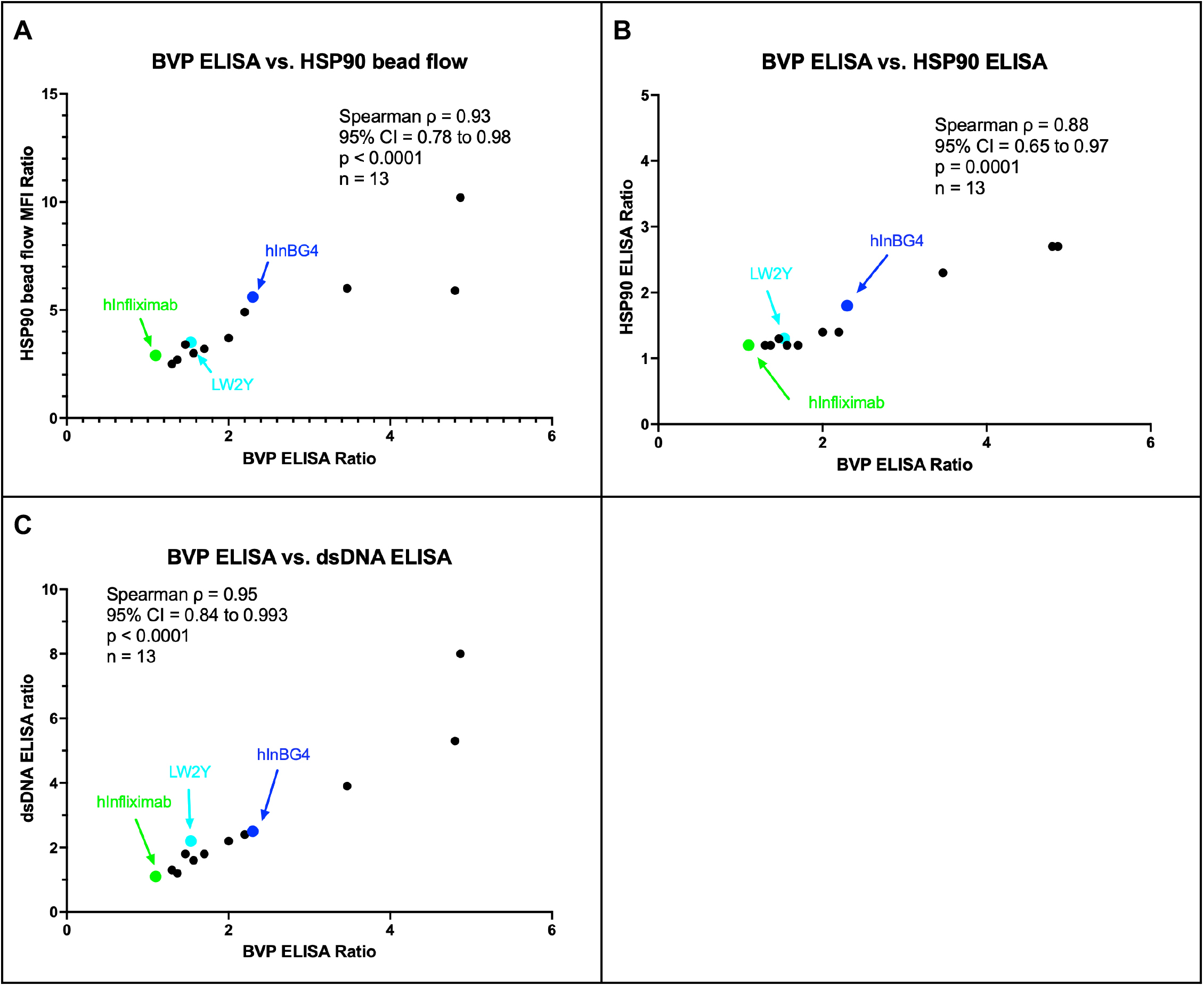
Orthogonal polyspecificity assays of single amino acid mutant clones of hInBG4 (purified IgG n = 13): Strong concordance was observed between BVP ELISA ratio and other assays (HSP90 ELISA ratio, dsDNA ELISA ratio, and HSP90 bead flow)

**Figure 4S.**
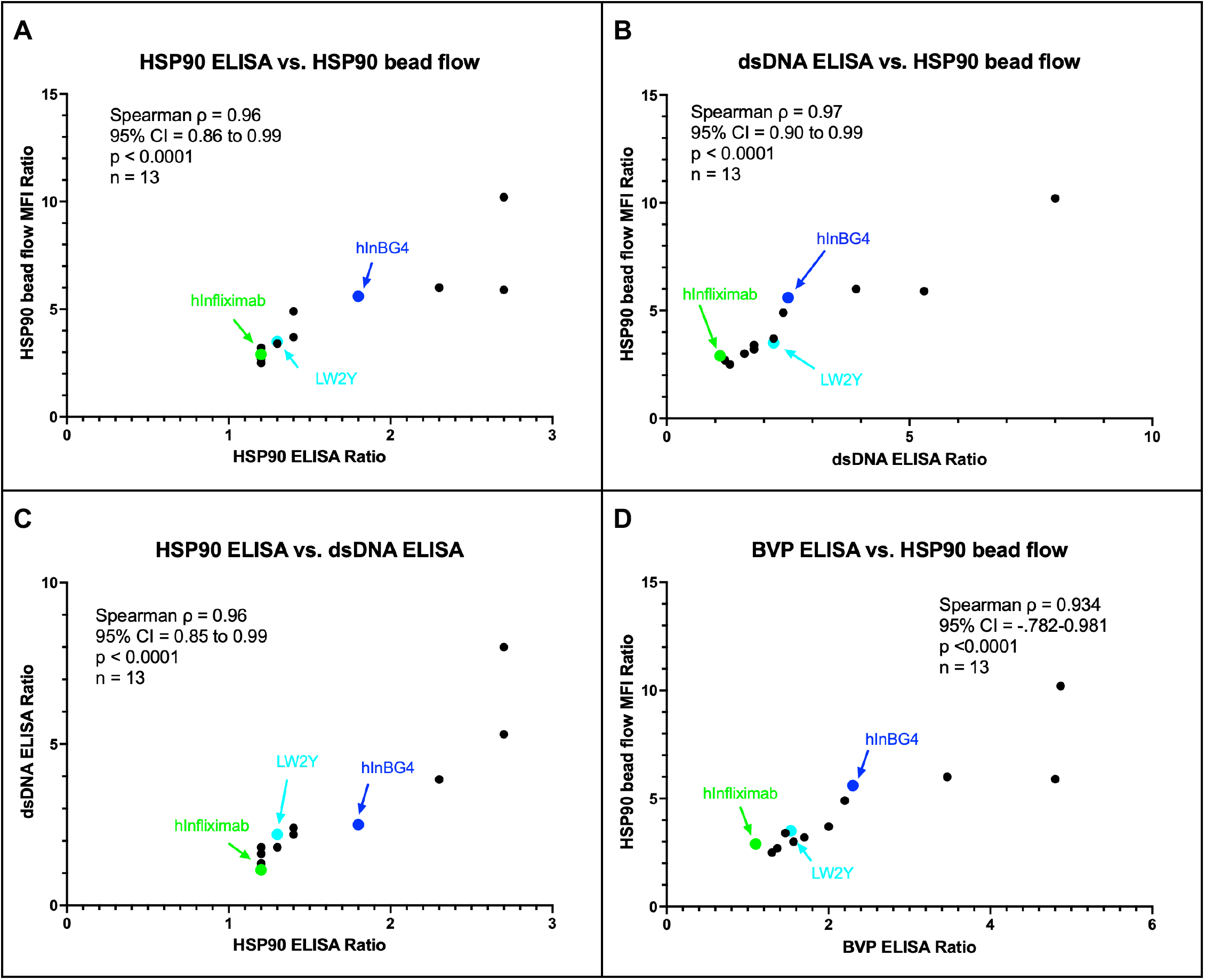
Orthogonal polyspecificity assays of single amino acid mutant clones of hInBG4 (purified IgG n = 13): Strong concordance was observed among HSP90 ELISA, dsDNA ELISA, and HSP90 bead flow.

**Table 1S.**
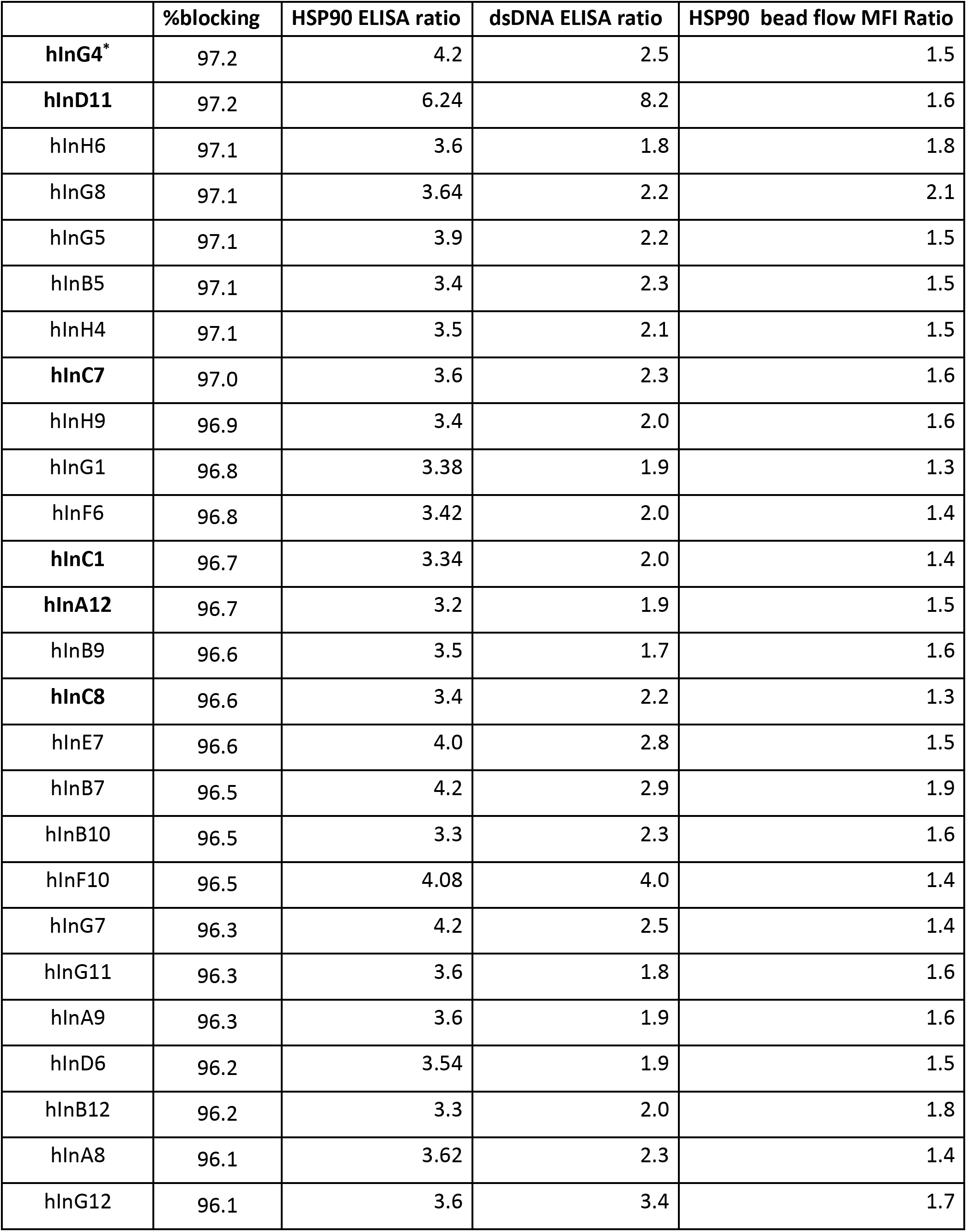
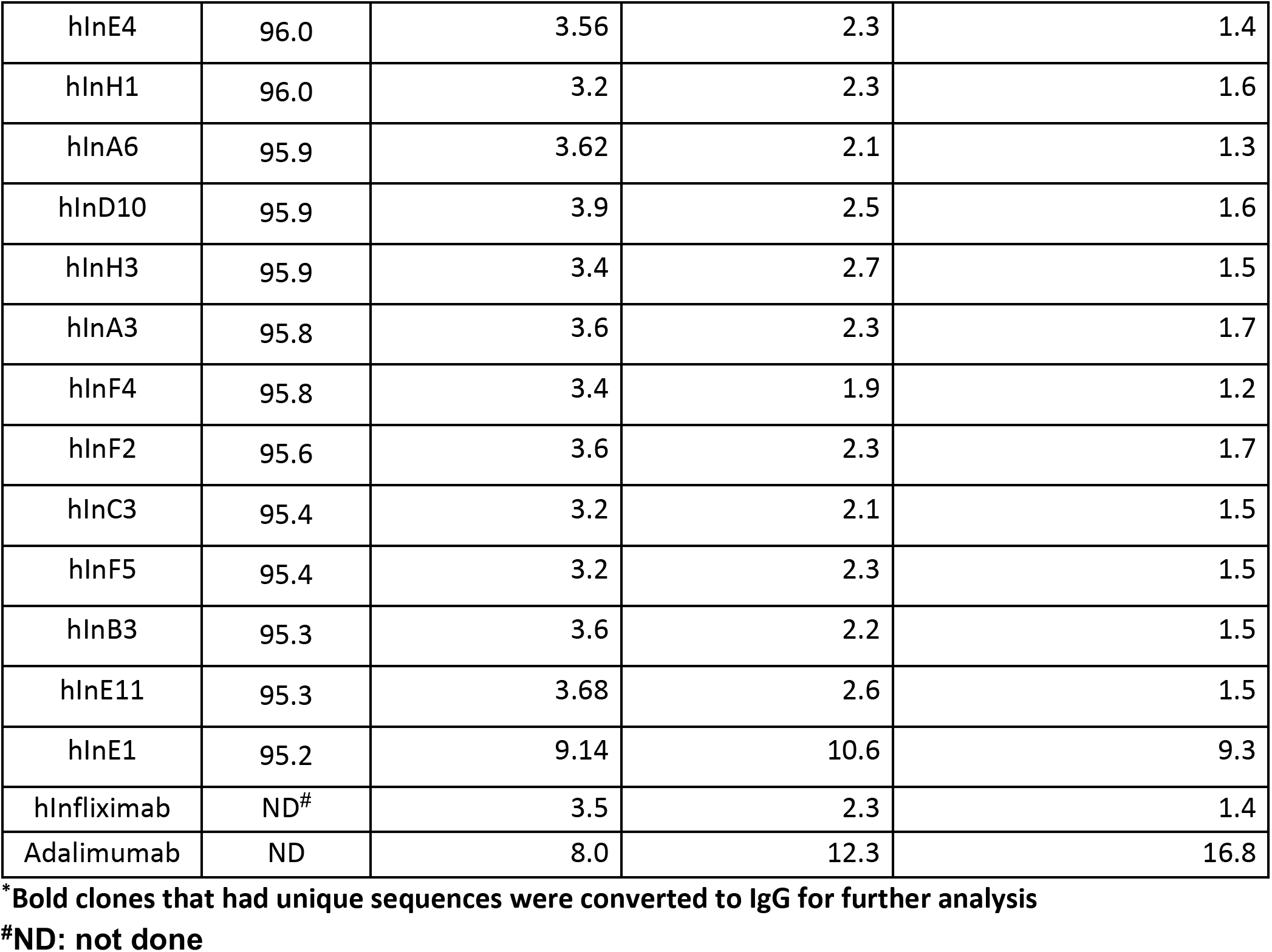
Crude Fab %blocking and PSR data of STEM™-engineered clones.

## REFERENCES

1. Tracey D, Klareskog L, Sasso EH, Salfeld JG, Tak PP. Tumor necrosis factor antagonist mechanisms of action: a comprehensive review. Pharmacol Ther. 2008;117:244–279.

2. Baert F, Noman M, Vermeire S, Van Assche G, D’Haens G, Carbonez A, Rutgeerts P. Influence of immunogenicity on the long-term efficacy of infliximab in Crohn’s disease. N Engl J Med. 2003;348:601–608.

3. Lerch TF, Sharpe P, Mayclin SJ, Edwards TE, Lee E, Conlon HD, et al. Infliximab crystal structures reveal insights into self-association. mAbs. 2017;9:874–883.

4. Jain T, Sun T, Durand S, Hall A, Houston NR, Hofer T, et al. Biophysical properties of the clinical-stage antibody landscape. Proc Natl Acad Sci U S A. 2017;114:944–949.

5. Fernández-Quintero ML, Ljungars A, Waibl F, Greiff V, Andersen JT, Gjølberg TT, et al. Assessing developability early in the discovery process for novel biologics. mAbs. 2023;15:2171248.

6. Bauer J, Rajagopal N, Gupta P, Gupta P, Nixon AE, Kumar S. How can we discover developable antibody-based biotherapeutics? Front Mol Biosci. 2023;10:1221626.

7. Sule SV, Dickinson CD, Lu J, Chow CK, Tessier PM. Rapid analysis of antibody self-association in complex mixtures using immunogold conjugates. Mol Pharm. 2013;10:1322–1331.

8. Liu Y, Caffry I, Wu J, Geng SB, Jain T, Sun T, et al. High-throughput screening for developability during early-stage antibody discovery using self-interaction nanoparticle spectroscopy. mAbs. 2014;6:483–492.

9. Wu J, Schultz JS, Weiss WF, Roberts CJ, Da Silva NA, Tessier PM. Discovery of highly soluble antibodies prior to purification using affinity-capture self-interaction nanoparticle spectroscopy. Protein Eng Des Sel. 2015;28:403–414.

10. Makowski EK, Wu L, Gupta P, Tessier PM. Discovery-stage identification of drug-like antibodies using emerging experimental and computational methods. mAbs. 2021;13:1895540.

11. Hötzel I, Theil FP, Bernstein LJ, Prabhu S, Deng R, Quintana L, et al. A strategy for risk mitigation of antibodies with fast clearance. mAbs. 2012;4:753–760.

12. Xu Y, Roach W, Sun T, Jain T, Prinz B, Yu TY, et al. Addressing polyspecificity of antibodies selected from an in vitro yeast presentation system: a FACS-based, high-throughput selection and analytical tool. Protein Eng Des Sel. 2013;26:663–670.

13. Rabia LA, Zhang Y, Ludwig SD, Julian MC, Tessier PM. Net charge of antibody complementarity-determining regions is a key predictor of specificity. Protein Eng Des Sel. 2018;31:409–418.

14. Datta-Mannan A, Lu J, Witcher DR, Leung D, Tang Y, Wroblewski VJ. The interplay of non-specific binding, target-mediated clearance and FcRn interactions on the pharmacokinetics of humanized antibodies. mAbs. 2015;7:1084–1093.

15. Starr CG, Tessier PM. Selecting and engineering monoclonal antibodies with drug-like specificity. Curr Opin Biotechnol. 2019;60:119–127.

16. Igawa T, Ishii S, Tachibana T, et al. Antibody recycling by engineered pH-dependent antigen binding improves the duration of antigen neutralization. Nat Biotechnol. 2010;28:1203–1207.

17. Watkins JM, Watkins JD. An engineered monovalent anti–TNF-α antibody with pH-sensitive binding abrogates immunogenicity in mice following a single intravenous dose. J Immunol. 2022;209:829–839.

18. Hong S-T, Su Y-C, Wang Y-J, Cheng T-L, Wang Y-T. Anti-TNF alpha antibody Humira with pH-dependent binding characteristics: a constant-pH molecular dynamics, Gaussian accelerated molecular dynamics, and in vitro study. Biomolecules. 2021;11:334.

19. Entzminger KC, Fleming JK, Entzminger PD, Espinosa LY, Samadi A, Hiramoto Y, et al. Rapid engineering of SARS-CoV-2 therapeutic antibodies to increase breadth of neutralization including BQ.1.1, CA.3.1, CH.1.1, XBB.1.16, and XBB.1.5. Antib Ther. 2023;6:108–118.

20. Callaway HM, Hastie KM, Schendel SL, Li H, Yu X, Shek J, et al. Bivalent intra-spike binding provides durability against emergent Omicron lineages: results from a global consortium. Cell Rep. 2023;42:112014.

21. Entzminger KC, Johnson JL, Hyun J, et al. Increased Fab thermoresistance via VH-targeted directed evolution. Protein Eng Des Sel. 2015;28:365–377.

22. Wu SJ, Luo J, O’Neil KT, Kang J, Lacy ER, Canziani G, et al. Structure-based engineering of a monoclonal antibody for improved solubility. Protein Eng Des Sel. 2010;23:643–651.

23. Kelly RL, Le D, Zhao J, Wittrup KD. Reduction of nonspecificity motifs in synthetic antibody libraries. J Mol Biol. 2018;430:119–130.

24. Raybould MIJ, Marks C, Krawczyk K, Taddese B, Nowak J, Lewis AP, et al. Five computational developability guidelines for therapeutic antibody profiling. Proc Natl Acad Sci U S A. 2019;116:4025–4030.

25. Kabat EA, Wu TT, Perry HM, Gottesman KS, Foeller C. Sequences of Proteins of Immunological Interest. 5th ed. Bethesda, MD: National Institutes of Health; 1991. NIH Publication No. 91–3242.

26. Datta-Mannan A, Thangaraju A, Leung D, Tang Y, Witcher DR, Lu J, Wroblewski VJ. Balancing charge in the complementarity-determining regions of humanized mAbs without affecting pI reduces non-specific binding and improves the pharmacokinetics. mAbs. 2015;7:483–493.

27. Roopenian DC, Akilesh S. FcRn: the neonatal Fc receptor comes of age. Nat Rev Immunol. 2007;7:715–725.

